# Adaptation of artificial diets for *Helicoverpa armigera* (Hübner) (Lepidoptera: Noctuidae) at laboratory rearing

**DOI:** 10.1101/396242

**Authors:** Caio C. Truzi, Hurian G. Holzhausen, José A. Chamessanga, Valéria L. Laurentis, Natalia F. Vieira, Alessandra M. Vacari, Sergio A. De Bortoli

**Author notes:** Corresponding author E-mail address (AMV). Author Contributions: conceptualization: AMV SADE methodology: AMV SADE formal analysis: AMV investigation: CCT HGH JAC VLL NFV writing (original draft preparation): CCT AMV SADE writing (review and editing): AMV SADE supervision: AMV SADE.

## Abstract

*Helicoverpa armigera* is an important pest of crops worldwide, and several studies have focused on the development of artificial diets for this species. However, studies evaluating the insect´s performance at nutritionally different diets are scarce. Larval development is dependent on the ratio of protein and carbohydrates. The aim of this study was to evaluate the biology and to compare the consumption and use of food of *H. armigera* larvae on diets with different protein levels. The nutritional index, the relative consumption rate, the relative metabolic rate, the relative growth rate, and the apparent digestibility were higher in the diet with added protein. On the other hand, the conversion efficiency of digested food was lower, resulting in a higher metabolic cost. In terms of biological aspects, larval survival was higher for the diet with average protein content and lower for the diet with a high protein level. The pupal period was longer for the diet with a higher protein content, while pupal survival was lower. Among the evaluated diets, the diet with an average protein content resulted in a higher net reproductive rate, a shorter time for the population to double in number, and the highest rates of population growth. The results suggest that lower or higher protein contents in the diets of *H. armigera* negatively affect the biological aspects of this species.

## Introduction

*Helicoverpa armigera* (Hübner) (Lepidoptera: Noctuidae) is an important pest of agricultural crops worldwide and stands out with polyphagia, facultative diapause, a high dispersal capacity, adaptation to different environments, and a high reproductive potential [1, 2]. This pest species can feed on vegetative organs and reproductive structures, causing significant losses and control costs of about US$ 5 billion worldwide [3, 4].

Although several studies have investigated artificial diets for *H. armigera*, few studies have been carried out evaluating its performance on nutritionally different diets. Larval development is dependent on the ratio of proteins and carbohydrates [5]. Protein digestion during the larval period is a complex process and performed by the proteases present in the insect’s gut [6].

In other lepidopteran species, such as *Plodia interpunctella* (Hübner) (Lepidoptera: Pyralidae), a diet low in proteins and carbohydrates can influence several biological aspects [7].

The development of artificial diets suitable for insects has allowed a great advance in integrated pest management programs. The consumption and use of food are basic conditions for insect growth, development, and reproduction, and the quantity and the proportion of nutrients in the food substrate of the larval stage influence the acceptance of the food, besides affecting the performance of adults, and may have more or less severe effects on the biology of insects, facilitating or impeding their development [8, 9, 10, 11, 12, 13].

In this context, the aim of this study was to evaluate the biology and to compare the consumption and use of food in *H. armigera* larvae on diets with different protein levels.

## Material and Methods

The rearing of *H. armigera* and the experiments were conducted at the Laboratory of Biology and Insect Rearing (LBIR), Department of Plant Protection, São Paulo State University (UNESP), Jaboticabal, SP. The insects were kept under controlled laboratory conditions at a temperature of 25 ± 1°C, a relative humidity of 70 ± 10%, and a photoperiod of 12 hours light/12 hours dark.

### Rearing of *H. armigera*

The larvae were placed in Petri dishes (6 cm diameter × 2 cm height) containing an artificial diet described by Greene et al. (1976), with modifications, and monitored until the larvae reached the pupal stage. At the pupal stage, they were separated by sex and transferred to copula and oviposition cages of PVC (20 cm diameter × 20 cm height), lined with paper towels and supported on a plastic cover (23.5 cm diameter × 3 cm height) lined with paper towels, with the top closed with voile fabric fastened with elastic. Twenty couples were conditioned per cage and fed with a 10% honey-water mixture on a piece of soaked cotton packed inside a plastic top (3 cm diameter × 1.5 cm height). The eggs were removed and placed in plastic containers (25 × 15 × 12 cm) until hatching.

### Artificial diets

The artificial diets used in this study are described in Greene et al. [14], with modifications. Three formulations were used; one for rearing (D_1_) and two with variations in the amount of protein, with double (D_2_) and with half (D_3_). The compositions of the diets are shown in Table 1.

**Table 1.**
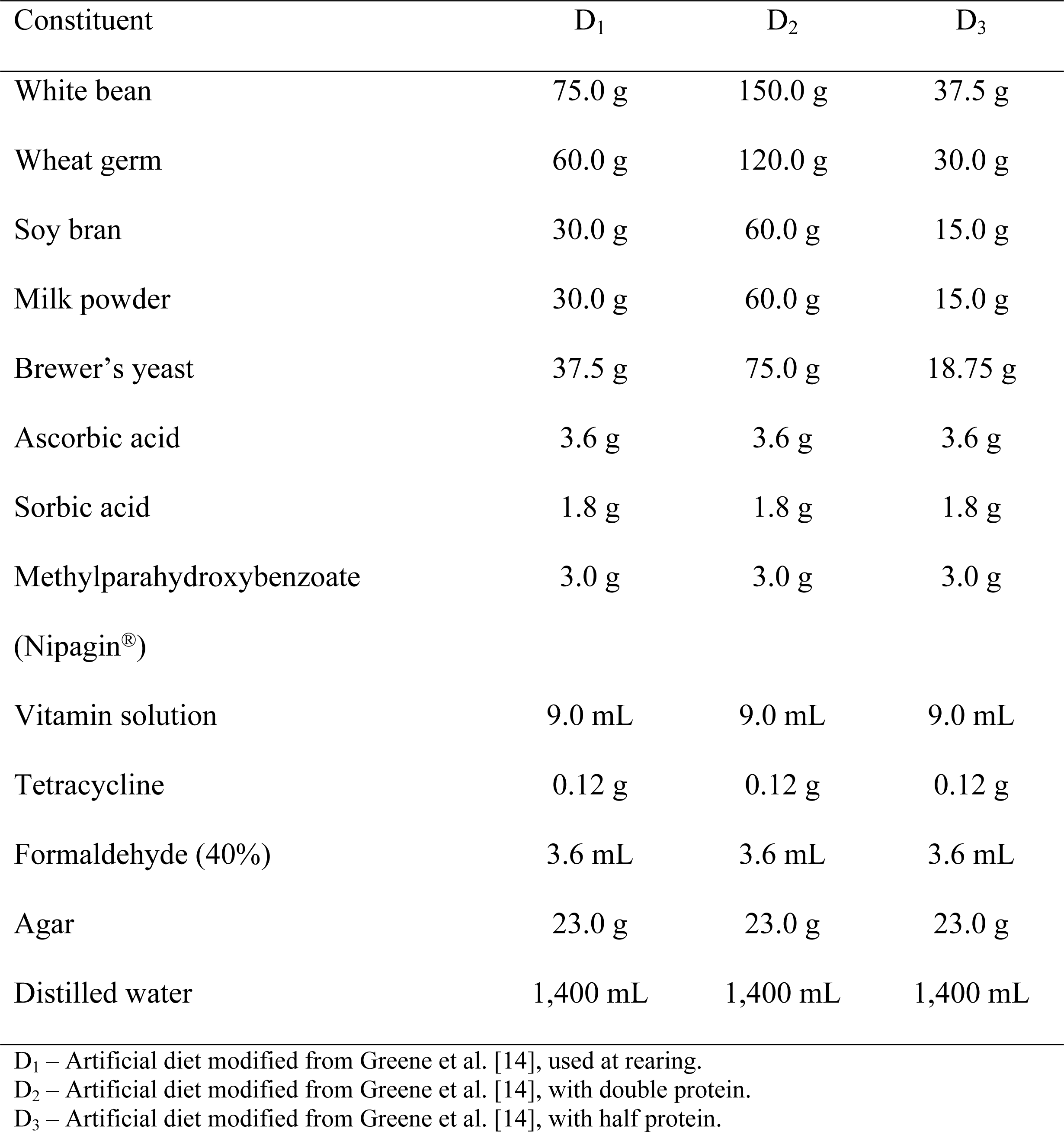
Composition of the artificial diets for *Helicoverpa armigera*.

**Table 2.** Composition of the vitamin solution used for artificial diets.

| Component | Amount |
| --- | --- |
| Niacinamide | 4.0 mg |
| Calcium pantothenate | 4.0 mg |
| Thiamine HCl | 1.0 mg |
| Riboflavin | 2.0 mg |
| Pyridoxine HCl | 1.0 mg |
| Folic acid | 1.0 mg |
| Biotin | 0.08 mg |
| Vitamin B12 | 0.008 mg |
| Distilled water | 400 mL |

### Nutritional indices

When the larvae reached the fourth instar, determined when they were about 12 mm in length [15] and had dark tubercles in the dorsal region of the first abdominal segment [16], 10 insects were removed from the Petri dishes, weighed, killed by freezing, and oven-dried. Another 10 insects were weighed and kept in the Petri dishes; where when they reached the fifth instar, 20 mm of length [15], confirmed by the presence of ecdysis, the same procedure as described above was performed. In addition, dietary leftovers and excrements from the insects during the fourth instar and 10 whole artificial diet cubes were weighed and oven-dried. After a drying period of 3 days, the diets and larvae were weighed. Thus, the weight of the dry larvae, the food consumed, and the weight gain of the larvae were obtained for the determination of the indices of food consumption and use.

The methodology proposed by Waldbauer [17] and modified by Scriber and Slansky Jr. [8] was adopted for determination the quantitative nutritional indices of fourth instar of the larvae stage. For the calculation of these indices, the following parameters were used (in dry matter weight):

T = duration of feeding period (days);

Af = weight of food supplied to the insect (g);

Ar = weight of leftover food provided to the insect (g) after T;

F = weight of excretory produced (g) during T;

B = (I – F) – M = weight gain by larvae (g) during T;

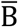 = mean weight of larvae (g) during T;

I = weight of food consumed (g) during T;

I – F = food assimilated (g) during T;

M = (I – F) – B = food metabolized during T (g).

The indices of consumption and use for each treatment were determined using the following equations:

Relative consumption rate 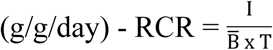;

Relative growth rate 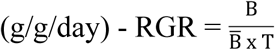;

Relative metabolic rate 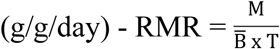;

Approximate digestibility 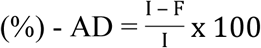;

Efficiency of conversion of ingested food 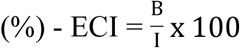;

Efficiency of conversion of digested food 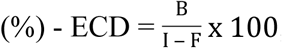;

Metabolic cost (%) - CM = 100 – ECD.

### Biological aspects

For each diet, 60 newly hatched larvae (< 24 hours) were individualized in Petri dishes (6 cm diameter × 2 cm height), where they were monitored until reaching the pupal stage. Artificial diet cubes (2 cm × 2 cm) were supplied and replaced after approximately 80% had been consumed. We evaluated the following biological parameters: larval period, larval survival, pupal weight at 24 hours, sex ratio, pupal period, and pupal survival.

After the emergence of adults, four cylindrical PCV cages (10 cm diameter × 20 cm height) were constructed for each diet, with the top closed with voile fabric fastened with elastic, supported on a plastic cover (15 cm diameter × 2 cm height) and lined with paper towels, where two couples of *H. armigera* that had emerged the same day were released. Adults were fed with a 10% honey-water mixture on a piece of soaked cotton packed inside a plastic top (3 cm diameter × 1.5 cm height). Based on daily observation, female fecundity (eggs/female) and longevity of male and female adults were determined.

With the biological parameters obtained, the parameters for the construction of fertility life tables were estimated, according to Birch [18], Price [19], and Southwood [20]: x = age of parental females, age is considered starting in the egg phase; lx = life expectancy to age x, expressed as a fraction per female; mx = specific fertility or number of offspring produced per female at age x; lx.mx = total number of females at age x. The growth parameters resulting from life tables were R_0_ = net rate of population growth, T = mean generation time, r_m_ = intrinsic rate of increase, λ = finite rate of increase. In addition to these parameters, we also determined Dt, which is the time required for the population to double in number, according to Krebs [21].

The growth parameters (R_0,_ T, r_m_, λ e Dt) were calculated using the following equations:

R_0_ = Σ(lx.mx);

T = Σ(x.lx.mx)/Σ(lx.mx);

r_m_ = ln R_0_/T;

λ = e^rm^;

Dt = ln(2)/r_m_.

### Statistical analysis

Data from nutritional indices and biological characteristics of adult *H. armigera* specimens on different diets were subjected to Kolmogorov and Bartlett tests to determine normality and homogeneity of variance. Data from the RCR, RGR, RMR, ECD, weight of the fresh and dry larvae, and pupal period were transformed by the root of x + 0.5 to meet the requirements of the analysis of variance (ANOVA) and then analyzed using PROC ANOVA. Means were compared by Tukey’s test (P < 0.05). Data for larval period, larval survival, weight of pupae, pupal period, pupal survival, fecundity of females, and longevity of males and females were compared using the Student Newman Keuls test. All analyses were conducted using the software package SAS [22].

Population parameters of fertility life tables were estimated according to the Jackknife methods for estimating the confidence intervals of the parameters and for allowing comparisons between treatments, as described by Maia et al. [23]. Estimates were conducted using SAS software [22].

## Results

### Nutritional indices

The highest fresh weight of *H. armigera* in the fifth instar was obtained in D_3_, whereas in D_2_ the caterpillars had the lowest weight, with a variation greater than 60 mg. Regarding dry weight, all treatments were similar (Fig. 1).

**Fig 1.**
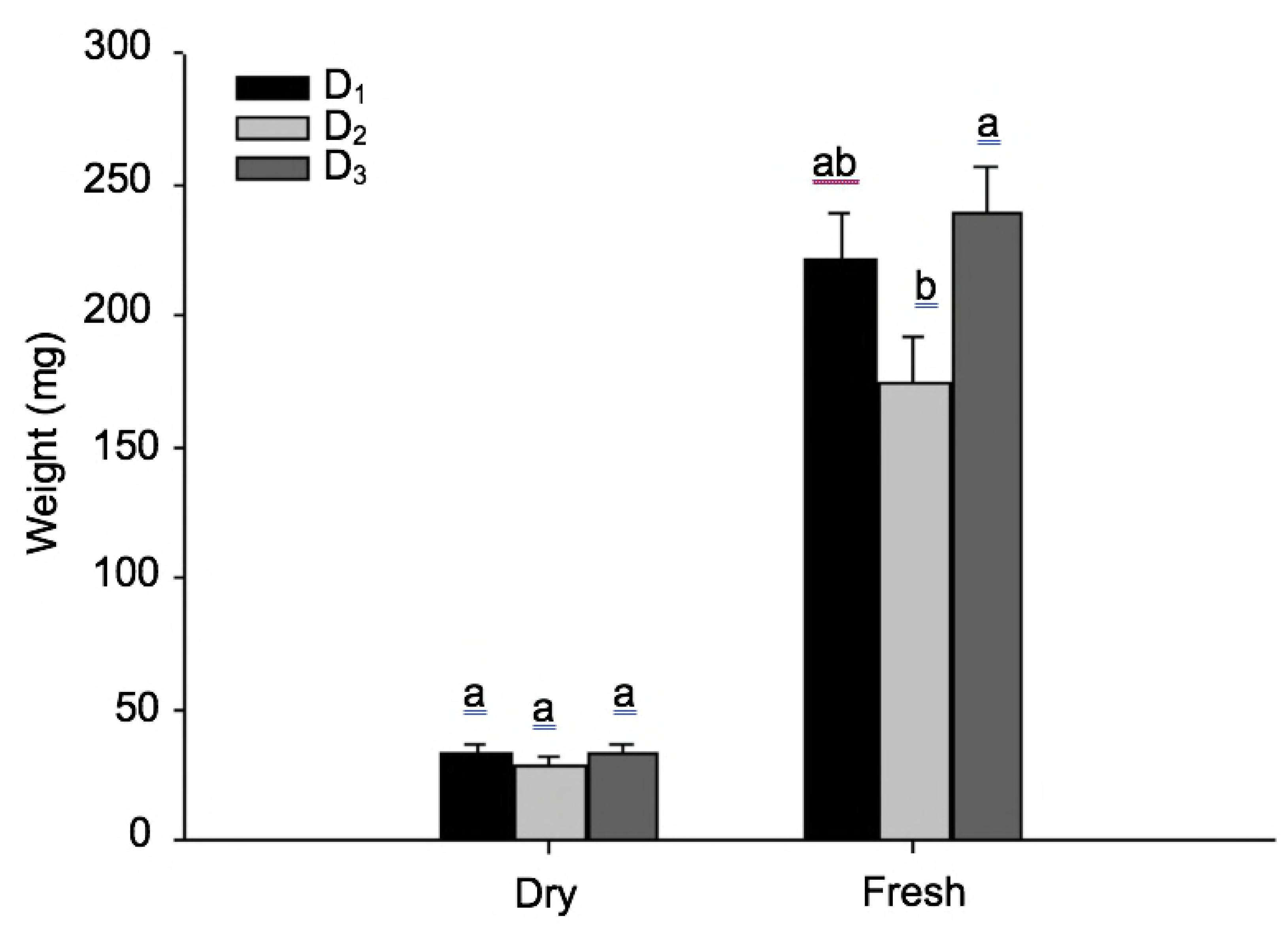
Dry and fresh weight of *Helicoverpa armigera* larvae on artificial diets. D_1_ – Artificial diet modified from Greene et al. [14], used at rearing. D_2_ – Artificial diet modified from Greene et al. [14], with double protein. D_3_ – Artificial diet modified from Greene et al. [14], with half protein.

Relative consumption rate (RCR), relative metabolic rate (RMR), and apparent digestibility (AD), which represents the percentage of the feed that was effectively assimilated, were higher in D_2_. For D_1_ and D_3_, these indices were smaller and similar to each other, differing in AD, which was lower for D_3_ (Table 3).

**Table 3.** Nutritional indices of *Helicoverpa armigera* on artificial diets.

| Index | Diet |  |  |
| --- | --- | --- | --- |
|  | D<sub>1</sub> | D<sub>2</sub> | D<sub>3</sub> |
| RCR (g/g/day) | 2.3 ± 0.40 b<sup>1</sup> | 5.5 ± 6.48 a | 1.9 ± 0.08 b |
| RGR (g/g/day) | 0.7 ± 0.00 b | 1.6 ± 0.41 a | 0.5 ± 0.01 b |
| RMR (g/g/day) | 0.7 ± 0.18 b | 2.5 ± 0.46 a | 0.3 ± 0.02 b |
| AD (%) | 58.4 ± 2.42 b | 73.2 ± 2.17 a | 45.8 ± 10.22 c |
| ECI (%) | 30.8 ± 1.90 a | 30.3 ± 1.21 a | 29.1 ± 2.21 a |
| ECD (%) | 54.4 ± 4.55 a | 41.9 ± 2.50 b | 63.8 ± 1.93 a |
| CM (%) | 45.6 ± 4.55 b | 58.1 ± 2.50 a | 36.2 ± 1.93 b |

<sup>1</sup>Means ± SE followed by the same letter on the line do not differ by the Tukey test (P > 0.05). D<sub>1</sub> – Artificial diet modified from Greene et al. [14], used at rearing. D<sub>2</sub> – Artificial diet modified from Greene et al. [14], with double protein. D<sub>3</sub> – Artificial diet modified from Greene et al. [14], with half protein.

Regarding the relative growth rate (RGR), the lowest values were for D_1_ and D_3_, while for the efficiency of conversion of ingested food (ECI) the values were similar for the three diets, varying from 29.1 to 30.8%. On the other hand, the efficiency of the conversion of digested food (ECD) was lower in D_2_, which led to a higher metabolic cost (CM) for this diet; while ECD was higher for D_1_ and D_3_ and CM was lower (Table 3).

### Biological characteristics

For the larval period, the insects fed with different diets did not present significant differences, with larval periods varying from 18.2 to 21.0 days. Survival in this period was higher for D_1_ and lower for D_2_, with a variation of 38.1%, while D_3_ presented intermediate survival (Table 4).

**Table 4.** Biological characteristics of the larval and pupal stages of *Helicoverpa armigera* on artificial diets.

| Characteristic | Diet |  |  |
| --- | --- | --- | --- |
|  | D<sub>1</sub> | D<sub>2</sub> | D<sub>3</sub> |
| Larval period (days) | 18.2 ± 0.53 a<sup>1</sup> | 20.9 ± 1.54 a | 21.0 ± 4.97 a |
| Larval survival (%) | 60.0 ± 7.29 a | 21.9 ± 4.63 b | 40.0 ± 7.29 ab |
| Pupae weight (mg) | 335.2 ± 10.23 a | 337.6 ± 12.29 a | 310.6 ± 9.08 a |
| Sex ratio | 0.4 ± 0.05 a | 0.4 ± 0.25 a | 0.3 ± 0.09 a |
| Pupal period (days) | 14.9 ± 0.36 b | 18.5 ± 1.06 a | 14.6 ± 0.21 b |
| Pupal survival (%) | 42.5 ± 5.00 a | 10.0 ± 4.68 b | 32.5 ± 6.37 a |

<sup>1</sup>Means ± SE followed by the same letter on the line do not differ by the Student Newman Keuls (P > 0.05). D<sub>1</sub> – Artificial diet modified from Greene et al. [14], used at rearing. D<sub>2</sub> – Artificial diet modified from Greene et al. [14], with double protein. D<sub>3</sub> – Artificial diet modified from Greene et al. [14], with half protein.

The weight of pupae at 24 hours of age and the sex ratio did not differ significantly between the diets evaluated, with the weight varied between 310.6 and 337.6 mg and the sex ratio between 0.3 and 0.4. The different diets influenced the pupal period of *H. armigera*, where the duration was higher for D_2_ and similar between D_1_ and D_3_ (Table 4).

Survival to the end of the pupal phase was lower for D_2_, while D_1_ and D_3_ showed similar survival rates (Table 4).

In adulthood, there was no significant difference in terms of longevity between males and females. However, females presented a higher longevity when compared to males in more than 3 days (Table 5).

**Table 5.** Biological characteristics of *Helicoverpa armigera* adults on artificial diets.

| Characteristic | Diet |  |  |
| --- | --- | --- | --- |
|  | D<sub>1</sub> | D<sub>2</sub> | D<sub>3</sub> |
| Male longevity (days) | 6.7 ± 1.96 a<sup>1</sup> | 5.5 ± 1.55 a | 5.0 ± 1.12 a |
| Female longevity (days) | 10.1 ± 1.84 a | 9.2 ± 1.31 a | 9.1 ± 1.33 a |
| Fecundity (eggs/female) | 229.7 ± 77.60 a | 268.7 ± 65.10 a | 206.5 ± 28.02 a |

<sup>1</sup>Means ± SE followed by the same letter on the line do not differ by the Student Newman Keuls (P > 0.05).
D<sub>1</sub> – Artificial diet modified from Greene et al. [14], used at rearing.
D<sub>2</sub> – Artificial diet modified from Greene et al. [14], with double protein.
D<sub>3</sub> – Artificial diet modified from Greene et al. [14], with half protein.

The different diets had no impact on female fecundity, which varied between 206.5 and 268.7 eggs/female (Table 5).

The Fertility Life Table, based on the results obtained for the biological parameters of *H. armigera*, showed differences between the evaluated diets. The net reproduction rate (R_0_), although higher for D_1_, was similar for all treatments, ranging from 22.7 to 61.3 females/females per generation. Regarding the average generation time (T), the lowest value was found for D_1_, while D_3_ presented the highest value. The intrinsic increase rate (r_m_) was higher for D_1_ and lower for D_3_, from 0.052 females/female/day. The finite increase rate (λ) was also higher for D_1_, presenting similar values between D_2_ and D_3_. The time for the population to double in size (Dt) was significantly lower for D_1_; while for D_3_, it took 3 more days for the population to double in the number of individuals when the larvae were fed on these diets (Table 6).

**Table 6.** Parameters of the fertility life table of *Helicoverpa armigera* on artificial diets.

| Characteristics | Diets |  |  |
| --- | --- | --- | --- |
|  | D<sub>1</sub> | D<sub>2</sub> | D<sub>3</sub> |
| R<sub>0</sub> | 61.3 a<sup>1</sup><br>(-20.37-143.00) | 23.7 a<br>(-108.66-156.05) | 22.7 a<br>(-4.65-50.11) |
| r<sub>m</sub> | 0.139 a<br>(0.1044-0.1756) | 0.091 ab<br>(-0.0791-0.2605) | 0.087 b<br>(0.0557-0.1180) |
| λ | 1.146 a<br>(1.1045-1.1866) | 1.096 b<br>(0.9123-1.2816) | 1.089 b<br>(1.0554-1.1233) |
| T | 30.1 c<br>(25.01-35.31) | 36.0 b<br>(34.29-37.81) | 36.8 a<br>(35.89-37.70) |
| Dt | 4.9 b<br>(3.62-6.21) | 7.4 ab<br>(-9.45-24.28) | 7.9 a<br>(4.87-10.86) |

<sup>1</sup>Means (confidence interval) followed by the same letter on the line do not differ by the Student's t test (P > 0.05). R<sub>0</sub> = net reproduction rate, r<sub>m</sub> = intrinsic increase rate, λ = finite increase rate, T = average generation time, Dt = time for the population to double in size. D<sub>1</sub> – Artificial diet modified from Greene et al. [14], used at rearing. D<sub>2</sub> – Artificial diet modified from Greene et al. [14], with double protein. D<sub>3</sub> – Artificial diet modified from Greene et al. [14], with half protein.

Among the diets evaluated, D_1_ had the highest net reproduction rate, the shortest time for the population to double in number, and the highest rates of population growth (Table 6).

## Discussion

By comparing the consumption, feed use, and biology of *H. armigera* larvae in diets with different protein contents, significant differences were identified.

The highest fresh weight of fourth instar larvae was related to the protein content, since the lower the protein content of the diet, the greater the weight of the larvae. These results differ from those found by Sarate et al. [5], who fed larvae with several host plants and observed that diets rich in proteins and/or carbohydrates resulted in higher larvae weight and a shorter larvae period. However, the dry larvae mass was similar between the different diets.

The relative consumption rate (RGR), representing the amount of food ingested per unit weight of the insect per day, and the relative metabolic rate (RMR), representing the amount of food spent in metabolism per unit weight, were higher for the diet with a higher protein content, demonstrating that larvae need a greater amount of food to meet their nutritional needs due to the high amount of protein required for their development.

Apparent digestibility (AD), which represents the percentage of ingested food that is effectively assimilated by the insect, was also higher for the diet with a higher protein content and lower for the diet with a lower protein content, suggesting that the amount of food assimilated by the insect was associated with the protein level; therefore, in diets rich in protein, a higher food intake is necessary to satisfy the nutritional needs of the insect.

In terms of the relative growth rate (RGR), which indicates the biomass gain by the insect in relation to its weight, the lowest values were found for the diets with the lowest amounts of protein; however, regarding the efficiency of the conversion of ingested food (ECI), which represents the percentage of food ingested that is transformed into biomass, there was no difference between the three diets. The efficiency of the conversion of digested food (ECD) showed that for diets with high protein contents, a low conversion of the diet to biomass occurred. Due to this low efficiency, this diet presented a high metabolic cost (CM).

The protein present in the diet did not influence the length of the larval period (18.2 to 21.0 days). Survival during this period was low in the high-protein diet, demonstrating that high levels of this nutrient in the diet may impair *H. armigera* population growth. Hamed and Nadeem [25], who evaluated different artificial diets to rear *H. armigera* in the laboratory, found a larval period varying between 15.6 and 42.8 days, and the results of this study are within this range. Truzi et al. [24], using an artificial diet similar to D_1_, found a larval period of 14.6 days, a higher value than observed in this study.

The variation in protein content did not influence pupal weight and sex ratio. However, the pupal period was influenced in that the high protein diet prolonged this phase of the insect’s life cycle. For the diet based on cotton seed and water chestnut, similar values have been found previously [25]. However, for artificial diets, this period was shorter [24].

The survival of insects in the pupal phase was lower for the diet with a high protein content, in which was also the case for the larval period, indicating again the negative influence of high protein levels on the biology of this pest species.

The diet in the larval period did not interfere with the longevity of adults. However, females presented greater longevity than males, which has also been observed in other studies using different diets [15, 26, 24].

Female fecundity was not influenced by the diet during the larval period. However, the values found were low when compared to other studies, which reported values between 440 and more than 2,500 eggs/female on different diets during the larval phase [24, 27, 28].

In relation to the fertility life table, for the net reproduction rate (R_0_), Truzi et al. [24] found a higher value for the artificial diet, with 255 females per female in each generation. For soybean cultivars, the rate ranged from 16.0 to 270.0 females per female [29], while for tomato cultivars, it was between 7.8 and 186.9 females per females [30]. The average generation time (T) was influenced by the protein content, and at lower protein levels, the insects took longer to complete the cycle. The intrinsic rate of increase (r_m_) was also affected by the low protein level, with a smaller number of females per female being produced per day. When *H. armigera* larvae were reared on soybean cultivars, the values were similar, ranging between 0.084 and 0.114 females per female each day [29], but for artificial diets and for different host plants, larger values have been observed in previous studies [24, 27]. For the finite rate of increase (λ), low and high protein levels resulted in a lower number of females per female each day. *Helicoverpa armigera* presented the shortest time for the population to double in size (Dt) at D_1_ with 4.9 days, but this period was higher than that found by Truzi et al. [24], which was 3.6 days.

The influence of different protein levels on *H. armigera* development, population growth, consumption indices, and feed use was studied, and the information obtained in this study can be used to adapt diets or to develop new diets for mass rearing of this insect species. In addition, future studies should evaluate the influence of different protein levels on successive generations of *H. armigera*.

## Conclusion

The most suitable artificial diet for the mass rearing of *H. armigera* in the laboratory was D_1_, with lower or higher levels of protein affecting the biological aspects of this pest species.

## Acknowledgements

We are very thankful to Dr. Sergio Antonio De Bortoli for the opportunity to carry out this work in the Laboratory of Biology and Insect Rearing (LBIR) within the Insect Biology of Postgraduate Program in Agriculture (Agricultural Entomology) of the FCAV/Unesp.

## References

1. Fitt GP. The ecology of Heliothis in relation to agroecosystems. Annu Rev Entomol. 1989;34: 17–52.

2. CABI. Crop protection compendium – Helicoverpa armigera. 2015. Available: http://www.cabi.org/cpc/datasheet/26757. Accessed 12 Sep. 2017.

3. Lammers JW, Macleod A. Report of a pest risk analysis: Helicoverpa armigera (Hübner, 1808). Plant Protection Service and Central Science Laboratory, Netherlands. 2007. 18 pp.

4. Fathipour Y, Sedaratian A. Integrated management of Helicoverpa armigera in soybean cropping systems. In: El-Shemy H, editor. Soybean – pest resistance. Cairo: InTech; 2013. pp. 231–280.

5. Sarate PJ, Tamhane VA, Kotkar HM, Ratnakaran N, Susan N, Gupta VS, Giri AP. Developmental and digestive flexibilities in the midgut of a polyphagous pest, the cotton bollworm, Helicoverpa armigera. J Insect Sci. 2012;12: 1–16.

6. Kotkar HM, Sarate PJ, Tamhane VA, Gupta VS, Giri AP. Responses of midgut amylases of Helicoverpa armigera to feeding on various host plants. J Insect Physiol. 2009;55: 663–670.

7. Bouayad N, Rharrabe K, Ghailani N, Sayah F. Effects of different food commodities on larval development and alpha-amylase activity of Plodia interpunctella (Hübner) (Lepidoptera: Pyralidae). J Stored Prod Res. 2008;44(4): 373-378.

8. Scriber JM, Slansky Jr. F. The nutritional ecology of immature insects. Annu Rev Entomol. 1981;26: 183–211.

9. Behmer ST. Insect herbivore nutrient regulation. Annu Rev Entomol. 2009;54: 165–187.

10. Panizzi AR, Parra JRP. Bioecologia e nutrição de insetos – base para o manejo integrado de pragas. Brasília: Embrapa; 2009. 1169 pp.

11. Schowalter TD. Insect ecology: an ecosystem approach. San Diego: Elsevier; 2011. 650 pp.

12. Parra JRP. The evolution of artificial diets and their interactions in science and technology. In: Panizzi AR, Parra JRP, editors. Insect Bioecology and Nutrition for Integrated Pest Management. Boca Raton: CRC Press; 2012. pp. 51-92.

13. Cohen AC. Insect diets: science and technology. Boca Raton: CRC Press; 2015. 473 pp.

14. Greene GL, Leppla NC, Dickerson WA. Velvetbean caterpillar: a rearing procedure and artificial medium. J Econ Entomol. 1976;69(4): 487–488.

15. Ali A, Choudhury RA, Ahmad Z, Rahman F, Khan FR, Ahmad SK. Some biological characteristics of Helicoverpa armigera on chickpea. Tunis J Plant Prot. 2009;4(1): 99–106.

16. Matthews M. Heliothinae moths of Australia: a guide to pest bollworms and related noctuid groups. Melbourne: CSIRO; 1999. 320 pp.

17. Waldbauer GP. The consumption and utilization of food by insects. Adv in Insect Physiol. 1968;5: 229–288.

18. Birch LC. The intrinsic rate of natural increase of on insect population. J Anim Ecol. 1948;17(1): 15–26.

19. Price PW. Insect Ecology. New York: John Willey; 1984. 607 pp.

20. Southwood TRE. Ecological methods. London: Chapman and Hall; 1978. 524 pp.

21. Krebs CJ. Ecology: the experimental analysis of distribution and abundance. New York: Harper Collins College Publishers; 1994. 801 pp.

22. SAS Institute. SAS: users guide: statistics. Version 9.0. Cary, USA; 2002.

23. Maia AHN, Luiz AJB, Campanhola C. Statistical inference on associated fertility life parameters using Jackknife technique: computational aspects. J Econ Entomol. 2000;93(2): 511– 518.

24. Truzi CC, Vieira NF, Laurentis VL, Vacari AM, De Bortoli SA. Development and feeding behavior of Helicoverpa armigera (Hubner) (Lepidoptera: Noctuidae) on different sunflower genotypes under laboratory conditions. Arthropod-Plant Interac. 2017;11: 797–805.

25. Hamed M, Nadeem S. Rearing of Helicoverpa armigera (Hub.) on artificial diets in laboratory. Pak J Zool. 2008;40(6): 447–450.

26. Jha RK, Tuan SJ, Chi H, Tang LC. Life table and consumption capacity of corn earworm, Helicoverpa armigera, fed asparagus, Asparagus officinalis. J Insect Sci. 2014;14(34): 1–17.

27. Razmjou J, Naseri B, Hemati SA. Comparative performance of the cotton bollworm, Helicoverpa armigera (Hübner) (Lepidoptera: Noctuidae) on various host plants. J Pest Sci. 2013;87: 29–37.

28. Naseri B, Golparvar Z, Razmjou J, Golizadeh A. Age-stage, two-sex life table of Helicoverpa armigera (Lepidoptera: Noctuidae) on different bean cultivars. J Agr Sci Technol. 2014;16: 19– 32.

29. Soleimannejad S, Fathipour Y, Moharramipour S, Zalucki MP. Evaluation of potential resistance in seeds of different soybean cultivars to Helicoverpa armigera (Lepidoptera: Noctuidae) using demographic parameters and nutritional indices. J Econ Entomol. 2010;103(4): 1420–1430.

30. Safuraie-Parizi S, Fathipour Y, Talebi AA. Evaluation of tomato cultivars to Helicoverpa armigera using two-sex life table parameters in laboratory. J Asia Pac Entomol. 2014;17: 837– 844.

